# Innate immunity mediator STING modulates nascent DNA metabolism at stalled forks in human cells

**DOI:** 10.1101/2021.04.16.440118

**Authors:** Vy N. Nguyen, Salomé Brunon, Maria N. Pavlova, Pavlo Lazarchuk, Roya D. Sharifian, Julia M. Sidorova

## Abstract

The cGAS/STING pathway, part of the innate immune response to foreign DNA, is known to be activated by cell’s own DNA arising from the processing of the genome, including the excision of nascent DNA at arrested replication forks. We found STING activation to affect nascent DNA processing, suggesting a novel, unexpected feedback connection between the two events. Depletion of STING suppressed and re-expression of the protein in STING-deficient cells upregulated degradation of nascent DNA. Fork arrest was accompanied by the STING pathway activation, and a STING mutant that does not activate the pathway failed to upregulate nascent strand degradation. Consistent with this, cells expressing the STING mutant had a reduced level of RPA on parental and nascent DNA of arrested forks as well as a reduced CHK1 activation compared to the cells with wild type STING. Together our findings reveal a novel connection between replication stress and innate immunity.

## Introduction

Innate immunity is a universal cell-intrinsic mechanism of protection against bacteria and viruses. The cGAS DNA sensor and the mediator STING are a part of the innate immunity pathway that responds to foreign DNA in the cytoplasm by inducing pro-inflammatory cytokines and type I interferons (Hopfner & Hornung, 2020). The cGAS/STING cascade is also triggered by the fragments of cells’ own genomic DNA in the cytoplasm (Chen *et al*, 2016) generated during DNA damage and repair (Alvarado-Cruz *et al*, 2021; de Oliveira Mann & Kranzusch, 2017; Harding *et al*, 2017), or as a consequence of telomere metabolism (Chen *et al*, 2017; Nassour *et al*, 2019), cellular aging (Glück *et al*, 2017; Lan *et al*, 2019; Takahashi *et al*, 2018), and other processes. The cGAS/STING pathway activation contributes to sensitization of cancer cells to radio- and chemotherapy (Chen *et al*, 2020; Li & Chen, 2018; Parkes *et al*, 2017), and STING or cGAS are often mutated or silenced in cancer cells (de Queiroz *et al*, 2018; Deschamps & Kalamvoki, 2017; Sun *et al*, 2013; Xia *et al*, 2016).

Stresses imposed on DNA replication, e.g. nucleotide depletion, result in slowing and stalling of replication forks. Stalled forks can undergo reversal and nucleolytic degradation of kilobases of nascent DNA via unwinding and resection of paired daughter strands. This phenomenon was first described in BRCA2-deficient cells treated with the ribonuclease inhibitor hydroxyurea, HU (Schlacher *et al*, 2011), and subsequently also demonstrated in BRCA-proficient cancer-derived cell lines and for several conditions of DNA damage or replication arrest (for review (Rickman & Smogorzewska, 2019)). Precise measurements of nascent DNA degradation were made possible by DNA fiber techniques (Quinet *et al*, 2017). In cancer cells, mutations causing upregulated nascent strand degradation (often referred to as a deficiency in the pathway of fork protection (Schlacher *et al*., 2011)) are associated with sensitivity to chemotherapies that interfere with DNA replication (Chaudhuri *et al*, 2016; Hill *et al*, 2018).

Replication fork arrest has been implicated in generating short extragenomic DNA species (Bétous *et al*, 2013; Shen *et al*, 2015; Yang *et al*, 2007). Furthermore, the nascent DNA removed from stalled forks via nucleolytic degradation can appear in the cytoplasm and induce the cGAS/STING pathway (Bhattacharya *et al*, 2017; Coquel *et al*, 2018). Fork protection/degradation is therefore one of the sources of immune-stimulatory self-DNA upstream of the cGAS/STING cascade.

We speculated that the cGAS/STING pathway, rather than being merely a downstream responder to fragmented genomic DNA, may provide a feedback that will modulate the very processes that trigger it. Here we asked if the status of STING in cells can have an import on fork protection. Our results indicate that STING depletion and re-expression respectively suppresses and enhances degradation of nascent DNA at stalled forks. The data are consistent with a positive feedback whereby nascent DNA degradation activates STING, which in turn enhances degradation.

## Results and Discussion

### Nascent strand degradation at stalled replication forks is affected by STING

The SV40-transformed fibroblast line GM639 displays moderate degradation of nascent DNA upon replication arrest by HU (Fig.1). This cell line and others also display slow incorporation of the 2nd label by forks restarting after HU (Sidorova *et al*, 2013). We depleted STING from GM639 (Fig.1A) to see if this affects the extent of degradation as measured by our DNA fiber assay (maRTA, (Sidorova *et al*, 2009)). Cells were pulse-labeled with two labels either consecutively or separated by a 5-6hr incubation with HU. Tracks of replication labeled with 1^st^ label (i.e. before HU) were measured in two-label tracks and in 1^st^ label only tracks (corresponding to, respectively, restarted and terminated forks), and compared to their equivalents in no HU samples (i.e. ongoing and terminated forks, Fig.1B). Thus, here the 1^st^ label, incorporated prior to HU, is the target for degradation in HU, as seen by the shortening of the 1^st^ label segments in restarted forks and 1^st^ label tracks of terminated forks compared to their no-HU counterparts. Addition of the 2^nd^ label distinguishes ongoing/restarted and terminated forks, which is important since the extent of degradation may vary between these two categories for biological as well as technical reasons.

STING depletion in GM639 fibroblasts partially suppressed the shortening of 1^st^ label tracks caused by HU, suggesting that the degradation of nascent DNA was reduced (Fig.1). The effect was more pronounced for the forks that did not restart within the first 30 min after HU (Fig.1C), but was also evident for the forks that restarted (Fig.1D). In contrast, slow restart and/or slow progression of post-HU forks, as evidenced by short tracks of the 2^nd^ label after HU, were not suppressed by STING depletion (Fig.1D, IdU).

**Figure 1.**
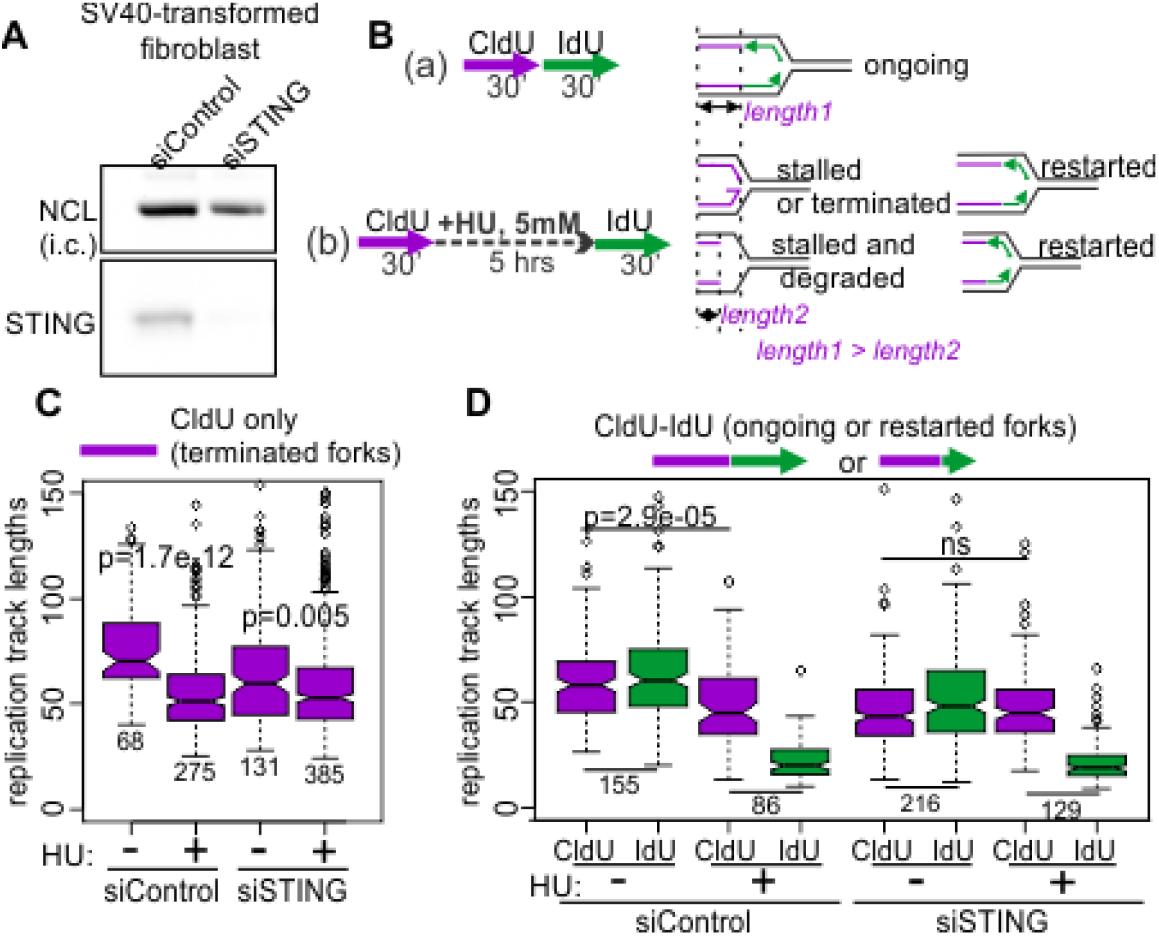
STING depletion affects nascent strand degradation at HU-stalled replication forks. A) A Western blot showing depletion of STING by siRNA in the SV40-transformed fibroblast line GM639. B) A maRTA experiment design to detect nascent strand degradation (left) and the categories of forks observed in untreated and HU-treated cells using this design (right). C) A boxplot of replication track length distributions comparing lengths of stalled or terminated forks (i.e. containing 1^st^ label only) in GM639 cells transfected with non-targeting and STING siRNAs. D) A boxplot of replication track length distributions comparing lengths of ongoing or restarted forks (i.e. containing 1^st^ and 2^nd^ labels) in GM639 cells transfected with non-targeting and STING siRNAs. (C) and (D) are representative of 3 independent experiments. Here and elsewhere the track length measurements are in pixels. 1 pixel approximately equals 1Kb. P values are derived in pairwise KS tests. Numbers of tracks measured are indicated beneath the boxplots.

To confirm these findings, we used U2OS cells which display moderate to no degradation of nascent DNA, depending on specific conditions (Thangavel *et al*, 2015). STING expression is silenced in U2OS (Deschamps & Kalamvoki, 2017). We stably re-expressed STING in U2OS cells from an integrated lentiviral construct pTRIP-SFFV-mtagBFP-2A STING (Cerboni *et al*, 2017) that enables selection of transduced cells by flow-sorting for BFP expression. STING expression was verified in cultured flow-sorted cells (Fig.2A), and cells were subjected to mRTA analyses using the same design as before (Fig.2B). Re-expression of STING augmented the HU-dependent shortening of tracks in both terminated and restarting forks (Fig.2C). Treatment with the MRE11 exonuclease inhibitor mirin during HU arrest and recovery reduced track shortening, confirming that exonucleolytic degradation of nascent DNA is a contributor to this phenotype (Extended Fig.2).

Depletion or re-expression of STING did not change fork progression under unperturbed conditions consistently across the cell lines (compare Fig.1D and 2C). Also, fork restart after HU was not affected by STING (not shown). Overall, the results argue that acute perturbation of STING can modulate the degree of nascent strand degradation exhibited by a given cell line: removal of STING ameliorates degradation, and supplementation of STING exacerbates it.

We were interested to see if STING depletion could also suppress the pathological nascent strand degradation associated with BRCA1 deficiency (Schlacher *et al*, 2012). We depleted 70-80% of STING (Fig.2D) in the BRCA1-negative UWB1.289 ovarian cancer cell line, and subjected the depleted cells and controls to the same regimen of labeling and HU arrest as in Fig.1 (Fig.2E). Due to BRCA1 deficiency, the tracks of forks that have been active prior to HU addition and then were stalled by HU (Fig.2E, lane 3) are considerably shorter compared to the tracks of forks that have not been subjected to HU arrest (Fig.2E lanes 1 and 2). When STING was depleted, the shortening of tracks subjected to HU arrest was largely suppressed (Fig. 2E, compare lanes 3 and 6). Taken together, the data suggest that STING can promote BRCA1-dependent and independent degradation of nascent DNA in at least some immortalized and cancer cells.

**Figure 2.**
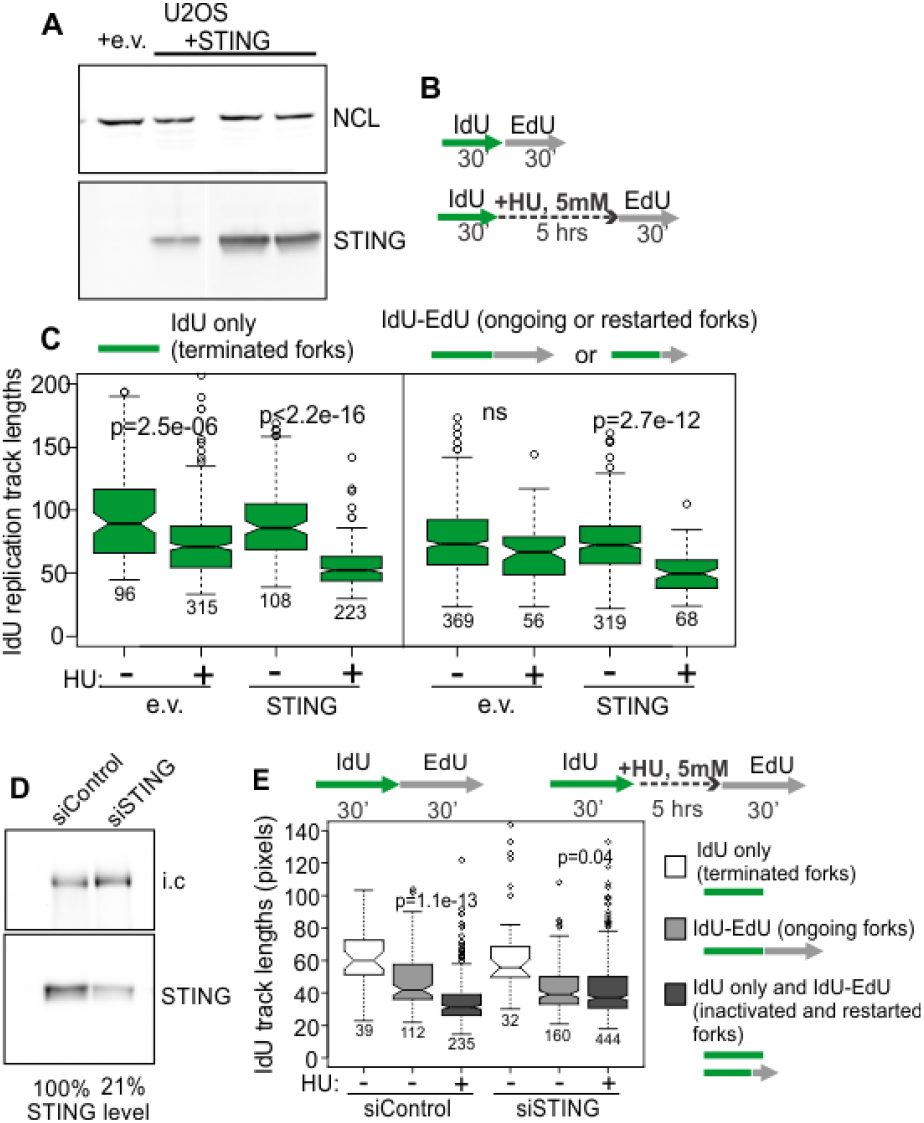
STING re-expression affects nascent strand degradation at HU-stalled replication forks. A) A Western blot of re-expression of STING in U2OS stably transfected with empty lentiviral vector pTRIP-SFFV-mtagBFP-2A (e.v.) or with the same vector expressing STING (pTRIP-SFFV-mtagBFP-2A STING). Cells were flow-sorted for BFP expression, and several fractions were cultured and analyzed for STING expression. The fractions with a higher expression of STING were used in maRTA experiments according to the design shown below. B) A maRTA experiment design to detect nascent strand degradation. C) Boxplots of replication track length distributions comparing lengths of stalled/terminated forks (left section) and ongoing/restarting forks (right section) in U2OS cells with empty vector or STING-expressing vector. The data represent 2 biological replicates. D) A Western blot of siRNA-mediated depletion of STING in BRCA1-deficient ovarian cancer line UWB1.289. i.c., internal control. E) An experimental design and a boxplot of 1^st^ label (IdU) track length distributions in terminated and ongoing forks (w/o HU, plotted separately), and in terminated or restarted forks (after HU, combined and plotted together since restarted forks are extremely rare at 30 min after HU in UWB1.289). The results represent two independent experiments.

### Effect on nascent strand degradation requires activation-competent STING

We next asked if the effect of STING on replication forks required activation of the STING pathway. This can be inferred from the induction of the pathway’s major transcriptional targets, type I interferons (IFNs) and inflammatory cytokines (Galluzzi *et al*, 2018). We detected induction of the classic targets of the pathway, the IFN beta, *IFNB1*, and interleukin 6, *IL6*, gene transcripts upon HU arrest conditions identical to those used for maRTA analyses. Induction of *IFNB1* and *IL6* mRNA by HU, albeit less dramatic than that achieved upon transfection of foreign DNA (e.g. interferon-stimulating DNA, ISD), was dependent on STING (Fig.3A,B). These data indicate that the pathway is indeed responsive to the HU-induced replication fork arrest.

To further investigate if activation of the STING pathway contributes to its effect at stalled replication forks, we introduced a S358A mutation into the STING expressed in U2OS cells. This mutation eliminates a STING phosphorylation site targeted by the TBK1 kinase and reduces binding of the transcription activator IRF3 to it, which is required for the activatory phosphorylation of IRF3 (Tanaka & Chen, 2012; Zhong *et al*, 2008). The STING S358A mutation thus markedly attenuates the human STING pathway activation in response to foreign DNA (Xie *et al*, 2018; Zhong *et al*., 2008).

Ectopic STING S358A was expressed at a level comparable to the wild type protein in our U2OS cells (Fig.3C). The STING 358A mutant failed to induce *IFNB1* mRNA in HU (Fig.3D), and as expected, did not support *IFNB1* or *IL6* mRNA activation upon interferon-stimulating DNA (ISD) transfection, confirming a defect in its ability to activate the downstream signaling cascade (Fig.3E,F). Interestingly, STING S358A also increased the basal level of *IL-6* mRNA (Fig.3F).

**Figure 3.**
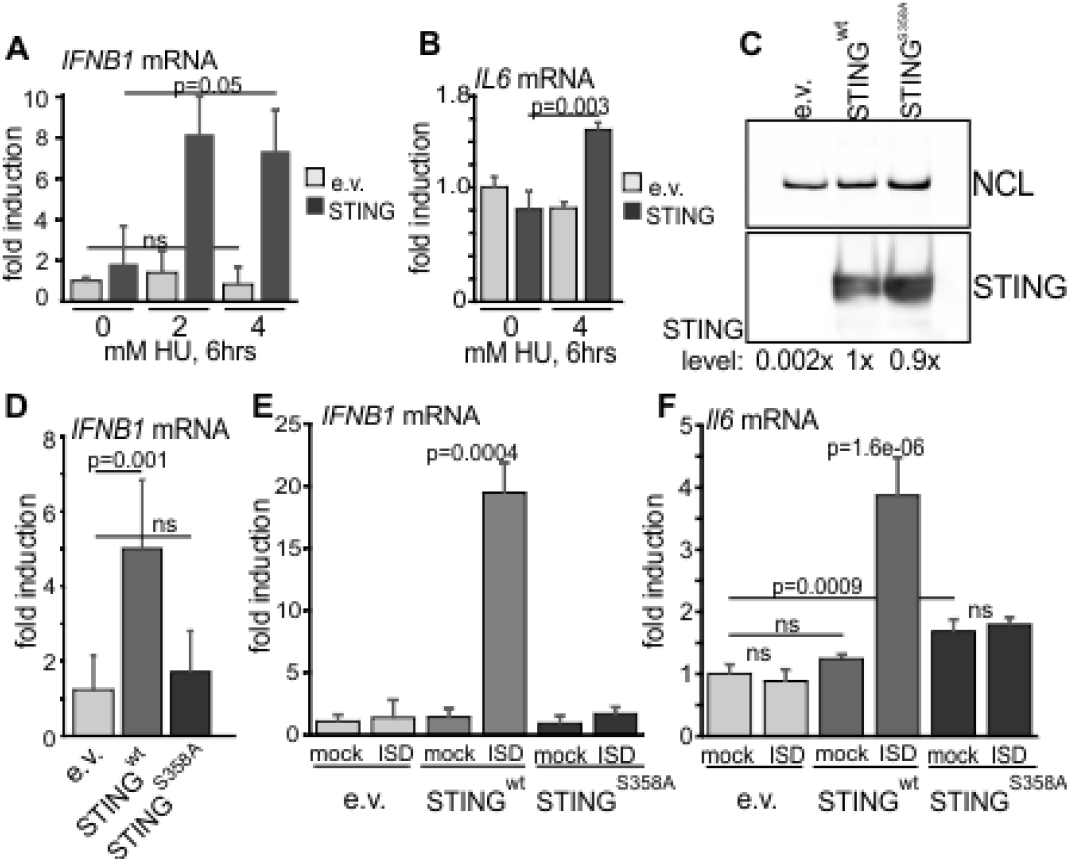
Serine 358 mutation of STING disrupts its ability to activate downstream transcriptional targets in response to exogenous DNA or replication arrest by HU. A) qPCR measurements of *IFNB1* mRNA induction in U2OS cells expressing STING or empty vector (e.v.) treated with 0, 2 or 4mM HU for 6 hrs. B) qPCR measurements of *IL6* mRNA induction in U2OS cells expressing STING or empty vector (e.v.) treated with 0 or 4mM HU for 6 hrs. A and B represent two independent experiments and P values were calculated on ΔΔCq values in one-tailed two-sample t-tests. C) A Western blot of U2OS cells expressing empty vector, wild type STING, or its S358A mutant. D) qPCR measurements of *IFNB1* maRNA induction in U2OS cells expressing the indicated STING variants or empty vector, incubated with 5mM hydroxyurea for 6 hrs. E-F) qPCR measurements of *IFNB1* (E) or *IL6* (F) mRNA induction in U2OS cells expressing the indicated STING variants or empty vector, and mock-transfected or transfected with interferon-stimulating DNA (ISD). qPCR results shown in D-F represent two independent experiments each.

Next, we performed mRTA analysis of nascent DNA degradation at HU-stalled forks. We used more than one labeling design to capture potentially different populations of forks while also being able to unambiguously identify the tracks that were generated by single forks. First, to increase the fraction of tracks formed by single, restarting forks, we increased recovery time to 60 min, with the 2^nd^ label present throughout this time. In these experiments, the STING-deficient control displayed no shortening of replication tracks at stalled forks (both terminated and restarting), whereas cells expressing the wild type STING markedly reduced track lengths (Fig.4A). In contrast, the track length phenotype of the STING S358A mutant was similar to the STING-deficient control (Fig.4A).

**Figure 4.**
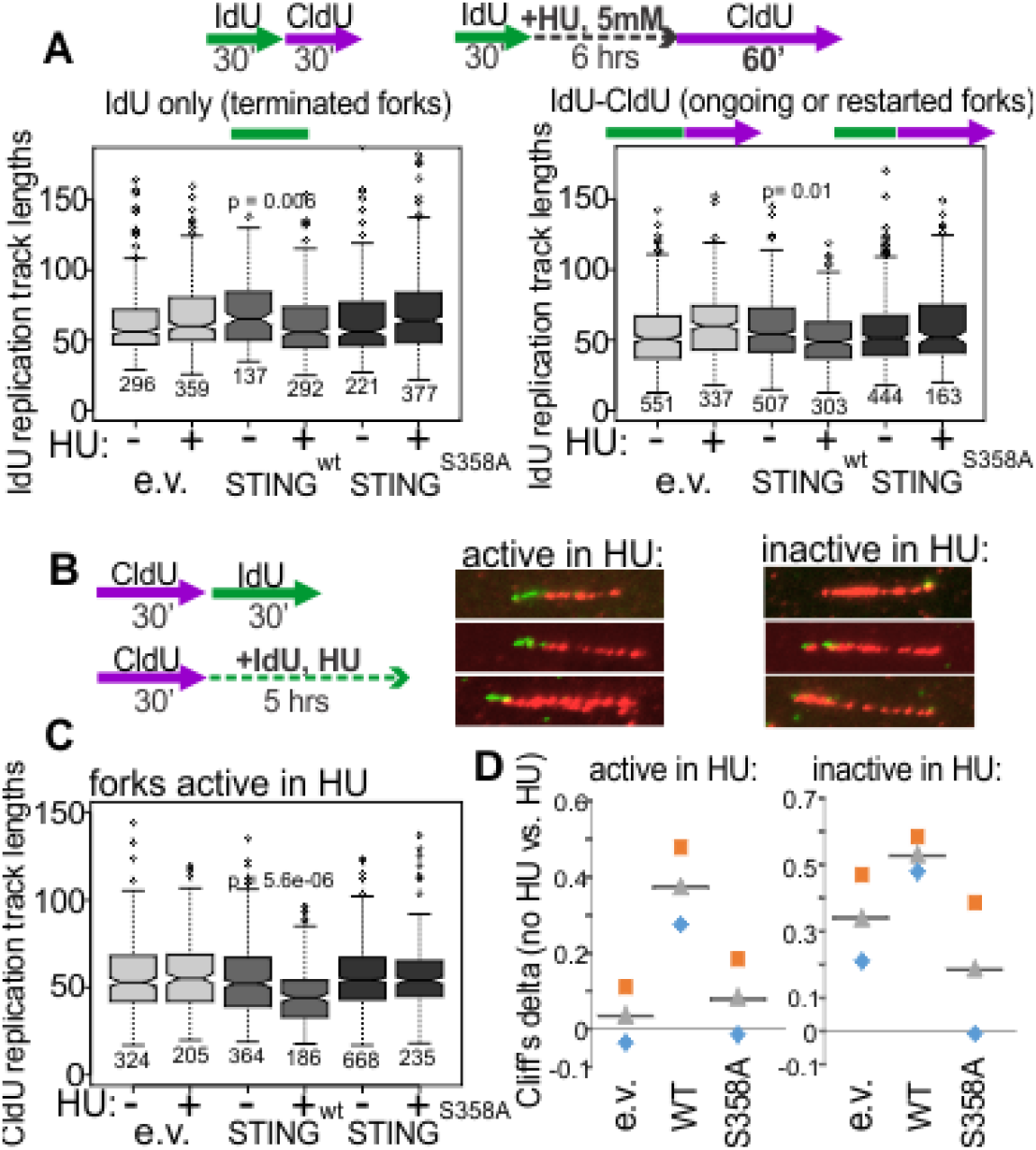
Activation of STING is important for its effect on degradation of nascent DNA. A) Experimental design and boxplots of 1^st^ label (IdU) track length distributions in terminated/stalled forks (left panel), or ongoing/restarted forks (right panel) in U2OS cells expressing the indicated transgenes. The results represent two independent experiments. P values are determined in KS tests. B) A labeling scheme and examples of forks that are active or inactive in the presence of 5mM HU. C) A boxplot of 1^st^ label (CldU) track length distributions in the active forks in U2OS cells expressing the indicated transgenes. The results represent two independent experiments. P values were determined in KS tests. D) Differences between 1^st^ label (CldU) track length distributions in untreated vs. HU-treated cells with the indicated transgenes were evaluated by calculating the respective Cliff’s delta values (see Materials and Methods for more detail). Here, positive values correspond to longer tracks in untreated cells compared to HU-treated cells. Cliff’s delta values from two independent experiments (squares and diamonds) and their averages (triangles) were plotted.

Next, we added the second label during the HU arrest rather than after it in order to distinguish between single forks and two or more converging/diverging forks regardless of whether they are able to restart after HU or not (Fig.4B). This design also allows to assess whether arrested forks experience a combination of nascent strand degradation and extension. We measured lengths of 1^st^ label tracks of two categories: tracks corresponding to single forks that incorporated at least a minimal measurable length of 2^nd^ label, approx. 15Kb (“active in HU”), and tracks that showed no evidence of extension activity, i.e. no 2^nd^ label incorporation (“inactive in HU”). STING presence correlated with a significant shortening of tracks in HU compared to the control for both categories, albeit the effect was more dramatic for the active forks than for the inactive forks (Fig.4C and 4D for Cliff’s deltas as metrics for the size of difference between track length distributions, see Materials and Methods for more detail). In contrast, STING S358A was very similar to the control when the active forks were measured, and interestingly, it was better than the control in suppressing the shortening of tracks of the inactive forks – an apparent gain-of-function phenotype (Fig.4D).

We also treated cells with a lowered dose of HU (1mM), under which 60-70% of forks, a percentage that typically accounts for all ongoing forks in a sample, were able to incorporate label during incubation with the drug. Under these conditions the presence of the wild type STING resulted in only a minor shortening of 1^st^ label segments that was not statistically significant compared to STING-negative control (data not shown). Together, the results rule out that the observed effects are due to an increase in nascent DNA fiber breakage either *in vivo* or *in vitro*, since the effect of STING is clearly observed in forks that show activity either during (Fig.4) or after HU arrest (Figs.1-2) and are captured in this state at point of analysis. The data also argue against a non-specific effect on DNA fiber lengths, since the STING-dependent impact on replication track lengths only manifests upon a near-complete arrest of forks that is most conducive to the cycles of reversal, degradation, and potentially, limited re-synthesis (Berti *et al*, 2020).

### STING affects the level of RPA at stalled forks and CHK1 phosphorylation in HU-arrested cells

The ssDNA-binding protein RPA regulates nascent strand degradation at more than one step (Berti *et al*., 2020; Bhat & Cortez, 2018; Duan & Pathania, 2020; Soniat *et al*, 2019). We hypothesized that RPA at stalled forks may be affected by the STING status as part of the mechanism that mediates STING’s effect on nascent strand degradation.

Total RPA levels assayed by Western blotting were similar between the STING-deficient, STING WT, and STING S358A U2OS (Extended Fig.5A). We next used Proximity Ligation Assay (PLA) and standard immunofluorescence (IF) in situ approaches to quantify RPA on nascent and parental ssDNA in S phase cells. PLA with antibodies against biotin (recognizing EdU-biotin conjugates) and the C terminus of the RPA subunit RPA32 (Fig.5A) was performed according to our described protocols (Lazarchuk *et al*, 2020; Lazarchuk *et al*, 2019). This PLA signal was specific to EdU-positive cells and detectable with and without HU (Fig.5A).

**Figure 5.**
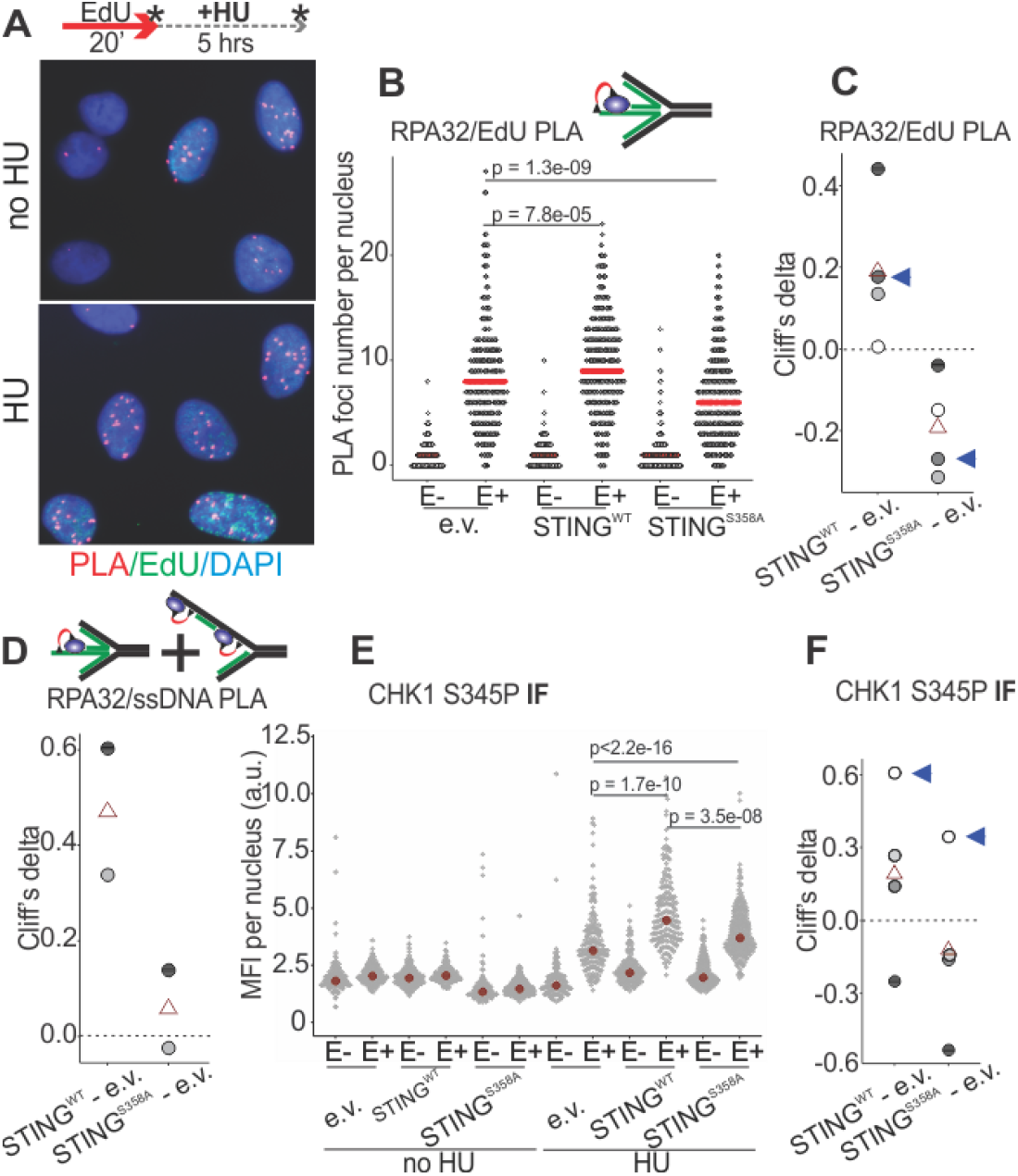
STING affects the levels of RPA on parental and nascent DNA and CHK1 activation in HU-treated cells. A) Examples of RPA32 to EdU PLA fluorescence in the control U2OS cells, treated as shown. Cells were labeled with EdU and harvested immediately or after 5hrs of HU arrest, as shown by asterisks. B) A schematic of the likely substrate for the RPA32/EdU PLA (red arc represents PLA signal) and representative distributions of RPA32/EdU PLA foci numbers in the U2OS cells with empty vector (e.v.) or the indicated STING transgenes. Cells were labeled as in (A) and EdU was Clicked to a mixture of biotin and Alexa488 azides at a molar ratio of 50:1 to enable simultaneous visualization of EdU-positive cells and PLA signal. PLA foci numbers are shown separately for EdU-negative (E-) and EdU-positive (E+) cells. Red lines are medians. P values were calculated in Wilcoxon tests. C) Magnitudes of differences between RPA32/EdU PLA foci distributions in STING WT vs. control and STING S358A vs. control cells were expressed as Cliff’s delta values and plotted. Shown are Cliff’s delta values for each of four independent experiments performed and quantified as in (B). Each experiment is identifiable by fill tone of the circle symbols. Triangle symbols are means. Blue arrowheads identify the values from the experiment shown in (B). D) Cliff’s delta values for two biological replicates of an experiment performed as in (B) and measuring RPA32/ssDNA PLA foci numbers. Likely substrates for the RPA32/ssDNA PLA are shown above the plot. E) The indicated U2OS cells were treated as in (A) and CHK1 S345P was visualized by IF in situ. MFI, mean fluorescence intensity, a.u., arbitrary units. F) Cliff’s delta values for MFI distribution differences between the indicated pairs were calculated for two independent experiments with biological replicates (i.e. four sets total) quantified as in (E). Values derived from each set are identifiable by fill tone. Blue arrowheads identify the values from the experiment shown in (E). For all Cliff’s deltas, positive values mean that the values of the first distribution in the comparison (e.g. STING WT in the STING WT – e.v. comparison) are overall higher than those of the second distribution, and negative values mean that the reverse is true.

Remarkably, HU-arrested wild type STING-expressing U2OS had an overall higher RPA32/EdU signal than the STING-deficient control (Fig.5B). In contrast, STING S358A-expressing cells consistently displayed less RPA32/EdU than the control (Fig.5B), which translated into a highly reproducible, dramatic differential between the wild type STING versus mutant STING-expressing cells (Fig.5C). This differential was also observed for the RPA32/ssDNA PLA signal (Fig.5D). Notably however, in this case mutant STING cells were comparable to the STING-deficient control rather than measuring below it (Fig.5D). Thus it is possible that STING S358A has a gain-of-function phenotype of reducing specifically the nascent DNA-bound RPA.

Phospho-RPA32 (S33P) measurements largely mirrored the above results. That is, in STING S358A cells the RPA32 S33P/EdU level (by PLA, Expanded Fig.5B,C) but not the total nuclear RPA32 S33P (by IF, Expanded Fig.5D) was below the STING-deficient control. At the same time, STING WT cells consistently displayed more of both RPA32 S33P/EdU and total RPA32 S33P than STING mutant cells (Expanded Fig.5B-D). Interestingly however, relative to the STING-deficient control, the RPA32 S33P/EdU signal was highly variable in STING WT cells (Expanded Fig.5C). One potential explanation may be an increased instability of phospho-RPA association with EdU-positive nascent DNA in STING WT cells.

When bound to the parental ssDNA at forks, RPA can promote stalled fork reversal, which is a prerequisite for nascent strand degradation (Berti *et al*., 2020; Bhat & Cortez, 2018), while nascent ssDNA-bound RPA participates in resection of this DNA (Duan & Pathania, 2020; Soniat *et al*., 2019). Therefore, detection of more RPA on parental and nascent ssDNA in the wild type STING-expressing cells (which can correspond to an enrichment per fork and/or to more forks in the RPA-enriched state) is consistent with their elevated nascent strand degradation. On the other hand, less RPA at ssDNA in the activation-incompetent STING S358A cells and a greater reduction of RPA specifically on the nascent DNA may suggest that fork reversal is reduced and the nascent ssDNA is less abundant or is unavailable for RPA binding and thus is protected from degradation. This nascent DNA phenotype is a gain-of-function by STING S358A over STING deficiency. It is tempting to speculate that this phenotype of STING 358A could be related to its gain-of-function hyper-protection of forks that do not incorporate any label in HU (compare Figs.4D and 5C).

At stalled forks, ssDNA-bound RPA is key to the activation of the ATR--CHK1 axis of the replication checkpoint (Bhat & Cortez, 2018). If in STING S358A cells the ssDNA-bound RPA is less abundant compared to the STING WT cells, we should expect an effect on the ATR-CHK1 checkpoint induction. Indeed, STING S358A cells consistently showed lower levels of the activated CHK1 kinase (CHK1 S345P) than STING WT cells (Fig.5E,F). Overall, the data reveal a specific pathway by which STING status can affect stalled fork metabolism, and establish a link between the replication checkpoint and innate immune responses to replication stress. However, further studies are needed to understand the pathway from STING to RPA recruitment and activity.

In summary, we showed that activation of STING during an HU-mediated replication fork arrest contributes to the regulation of degradation of nascent DNA at arrested forks. Since the excised nascent DNA can be a trigger for STING activation (Bhattacharya *et al*., 2017; Coquel *et al*., 2018), this may suggest that there is a positive feedback from the activated STING to fork processing. Such feedback may target specifically fork processing by modifying the activity of particular proteins, or have a broader effect on chromatin, with the upregulated nascent strand degradation being one of the consequences. In either scenario, altered loading of RPA onto parental and nascent DNA and the associated altered replication checkpoint induction, are involved. Arguing in favor of a targeted crosstalk between fork remodeling and the cGAS/STING pathway is the fact that the proteins MRE11 and SAMHD1, which facilitate nascent strand degradation, are also respectively a nucleic acid sensor and a negative regulator of the cGAS/STING pathway (Chen *et al*, 2018; Coquel *et al*., 2018; Kondo *et al*, 2013; Schlacher *et al*., 2011), though it is not clear how these, apparently dual, roles play out in the context of fork arrest and processing. Overall, our data are the first demonstration of a replication fork-associated phenotype for STING. While cGAS has been detected in the nucleus (Gentili *et al*, 2019; Liu *et al*, 2018; Volkman *et al*, 2019) and implicated in suppressing fork progression and nascent strand degradation (Chen *et al*., 2020), these functions are thus far considered to be STING-independent. Nevertheless, it is interesting to consider a possibility that STING may counteract cGAS at forks.

Fork protection and the cGAS/STING pathway are often suppressed in cancer cells (de Queiroz *et al*., 2018; Deschamps & Kalamvoki, 2017; Sun *et al*., 2013; Xia *et al*., 2016). Our data invite a possibility that loss of the cGAS/STING pathway can be advantageous vis-a-vis the chronic replication stress and increased nascent strand degradation of cancer cells by severing a positive feedback that amplifies degradation and checkpoint activation. In contrast, in normal cells where nascent strand degradation is minimal, STING activation may remain beneficial as it boosts the response to rare events of replication stress.

## Materials and Methods

### Cells and culture

The SV40-transformed human fibroblast GM639 (GM00639) has been described (Sidorova *et al*, 2008). The U2OS line was acquired from ATCC (ATCC HTB-96). UWB1.289 (DelloRusso *et al*, 2007) was a gift of Drs. Welcsh and Swisher. GM639 and U2OS were grown in Dulbecco Modified Minimal Essential Medium (DMEM) with L-glutamine, sodium pyruvate, 10% fetal bovine serum (Hyclone, Ogden, UT) and antibiotics. UW289.B1 was grown in 1:1 mixture of RPMI and MGEM (Lonza) with Single Quots (Lonza) with 3% FBS. All cell lines were kept in a humidified 5% CO2, 37°C incubator. Mycoplasma testing was performed regularly using the UW/FHCRC Cancer Consortium Shared Resource Specimen processing service \https://sharedresources.fredhutch.org/services/mycoplasma-testing.

### Drugs and other reagents

Stock of 5-iododeoxyuridine (IdU, Sigma-Aldrich) was at 2.5mM in PBS, 5-chlorodeoxyuridine (CldU, Sigma-Aldrich) was at 10mM in PBS, and 5-ethynyldeoxyuridine (EdU, Sigma-Aldrich or Click Chemistry Tools) was at 10mM in DMSO. IdU and CldU were used at a concentration of 50µM and EdU was used at 10 or 20µM. Hydroxyurea (Sigma-Aldrich) stock solution was at 1M in PBS and mirin (Calbiochem) was at 10mM in DMSO. All stocks were stored at -20°C.

### RNAi-mediated depletion

siRNAs against STING Hs_TMEM173_1 and Hs_TMEM173_4, and a Negative Control non-targeting siRNA were from Qiagen and transfected with lipofectamine RNAiMAX (Invitrogen) according to the manufacturer’s protocol. Experiments were performed 36 to 48 hrs post-transfection. Depletion was verified in each transfection by Western blotting.

### Constructs

pTRIP-SFFV-mtagBFP-2A STING and the parental empty vector were a gift from Nicolas Manel (respectively, Addgene plasmid # 102586; http://n2t.net/addgene:102586; RRID:Addgene_102586; and Addgene plasmid # 102585; http://n2t.net/addgene:102585; RRID:Addgene_102585). The S358A mutation was introduced into the STING ORF in this construct using Q5 site-directed mutagenesis kit (NEB). Virus generation from these constructs and cell transduction were as described (Sidorova *et al*., 2008). Live transduced cells were sorted on the Aria flow sorter based on the level of BFP expression.

### Antibodies

Antibodies were as follows: mouse α-biotin Cat. No. MB-9100 (Vector Laboratories); rat α-BrdU/CldU Cat. No. ab6326 (Abcam); mouse α-BrdU/IdU Cat. No.347580 (BD Biosciences); rabbit α-STING Cat. No.19851-1-AP (Proteintech), mouse α-NCL Cat. No. 396400 (Life Technologies), rabbit α-RPA32 Cat. No. A300-244A (Bethyl Labs), rabbit α-RPA32 S33P Cat. No. A300-246A (Bethyl Labs), rabbit CHK1 S345P Cat. No. 2348 (CST), mouse α-ssDNA MAB3034 (Millipore Sigma).

Proteins were visualized on Western blots by ECL (ThermoScientific) and quantified using FluorChem Imager (Alpha Inotech). For presentation, images were saved in TIFF format, adjusted for brightness/contrast and cropped in Adobe Photoshop, then assembled into figures in CorelDraw. Image brightness/contrast adjustments were made across all lanes of each protein measured. In some cases lane order was changed and extra lanes were deleted.

### Microfluidics assisted replication track analysis (maRTA)

This procedure was done as described (Kehrli & Sidorova, 2014; Sidorova *et al*., 2009). Microscopy of stretched DNAs was performed on the Zeiss Axiovert microscope with a 40x objective, and images were captured with the Zeiss AxioCam HRm camera. Lengths of tracks were measured in raw merged images using Zeiss AxioVision software. Fluorochromes were Alexa594 for CldU, Alexa488 for IdU, and Neutravidin Texas Red for EdU.

### Transfection of interferon-stimulating DNA (ISD)

2µg of ISD (Invivogen) was transfected into cells using lipofectamine 2000 (Invitrogen) per manufacturer’s protocol. Cells were incubated for 6hrs and harvested for RNA analyses. Mock-transfected controls received lipofectamine/Optimem mixture only.

### RNA isolation and qPCR

RNAs were isolated using RNeasy Plus RNA isolation kit (Qiagen). 2µg of RNA was reverse-transcribed using High Capacity cDNA Reverse Transcription kit (Applied Biosystems) per manufacturer’s protocol. cDNAs were diluted 1:10 and 1µl of diluted cDNA was used per qPCR reaction with iTaq Universal SYBR Green supermix (Bio-Rad) and the following pairs of primers: 5’ACAACTTTGGCATTGAA3’ and 5’GATGCAGGGATGATGTTCTG3’ for GAPDH; and Sigma-Aldrich KiCqStart predesigned pairs H_IFNB1_1 and H_IL6_1, Cat. No. KSPQ12012G for IFNb and IL6. Triplicate Ct values were averaged, normalized to GAPDH, and fold induction of mRNAs was determined according to the 2^−ΔΔCt^ method.

### Proximity Ligation Assay (PLA) and Immunofluorescence (IF) in situ

PLA was performed using DuoLink red detection kit and DuoLink anti-mouse and anti-rabbit antibodies (Millipore-Sigma Cat. No DUO92008, DUO92001, and DUO92002, respectively) as described previously (Lazarchuk *et al*., 2019), except that after formaldehyde fixation cells were washed in PBS, permeabilized by addition of 4°C 90% methanol in PBS and stored at -20°C prior to staining. Images of cells were collected under Zeiss Axiovert 200M microscope with 40X magnification objective using Micro Manager software. Digital images were analyzed with Fiji Image J software package with custom macros as described in (Lazarchuk *et al*., 2019) or with Cell Profiler software package.

### Statistical analysis

Statistical analyses and graphing of the data were done in R studio. P values for qPCR results were derived from pairwise t-tests on ΔΔCq values with Benjamini-Hochberg adjustment for multiple comparisons. P values for the rest of the assays were as follows. For continuous variables (e.g. track lengths, mean fluorescence intensities) p values were calculated in K.S. tests and for discrete variables (e.g. PLA foci) – in Wilcoxon tests. To quantify and concisely visualize pairwise differences between the distributions we calculated their Cliff’s deltas. In general, Cliff’s delta of distributions A vs. B can range from 1 (if all values in A are larger than all values in B) to -1 (if the reverse is true), and 0 value indicates that the distributions are completely overlapping.

## Acknowledgements

We are grateful to Dr. Kristin Eckert for critical reading of the manuscript. This work was supported by the NIH grant R01 GM115482 to J.S.

## Author contributions

MaRTA experiment design and data collection: V.N.N., J.S.; RT qPCR analyses for the cGAS/STING pathway induction: S.B. and J.S.; STING expression: V.N.N., J.S.; STING mutant construction: S.B. and M.P., siRNA-mediated STING depletion and Western blot analyses: R.S.; study design and manuscript writing: J.S.; PLA and IF analyses: P.L. and J.S.

## Expanded View Figures

**Expanded Figure 2.**
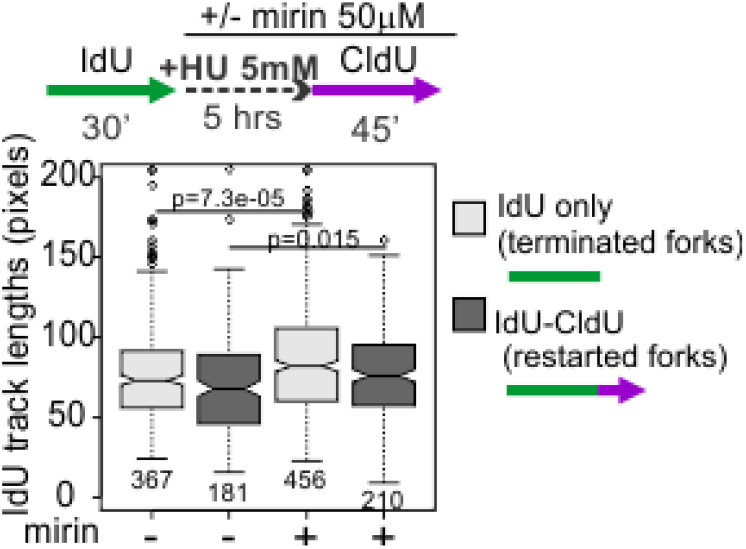
STING perturbations affect MRE11-dependent nascent strand degradation. An experimental design and a boxplot of 1^st^ label (IdU) track length distributions in terminated and restarting forks from U2OS expressing STING and treated or not treated with 50uM mirin during HU arrest and recovery.

**Expanded Figure 5.**
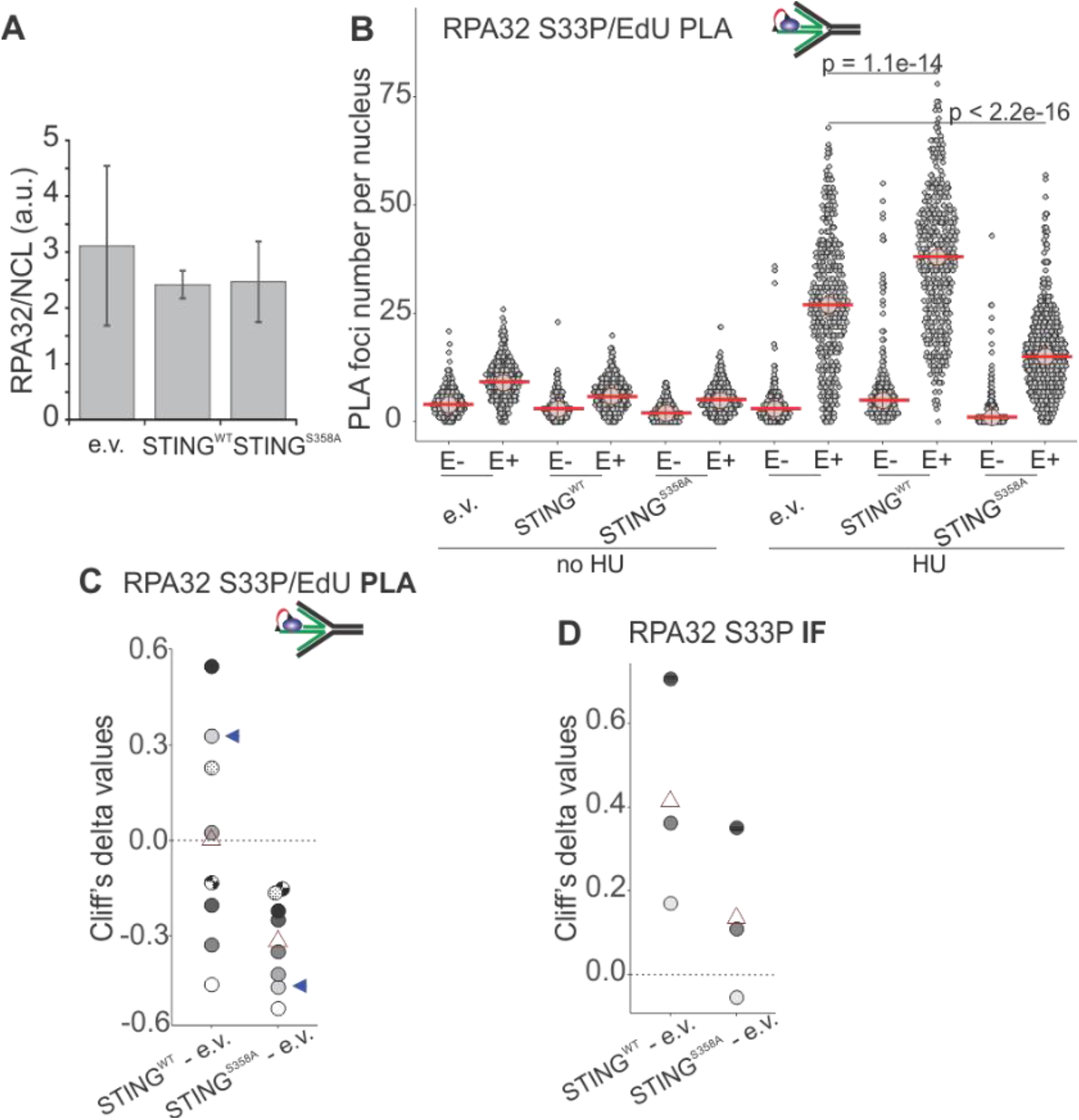
RPA32 S33P levels in HU-arrested cells are affected by the STING status. A) Quantification of Western blot results of RPA32 measurement in two biological replicate samples of U2OS cells expressing the empty vector (e.v.) or the indicated transgenes. B) Distributions of RPA32 S33P/EdU PLA foci numbers in the U2OS cells with empty vector or the indicated STING transgenes. Cells were labeled with EdU as in Fig. 5A and EdU was Clicked to a mixture of biotin and Alexa488 azides at a molar ratio of 50:1 to enable simultaneous visualization of EdU-positive cells and PLA signal. PLA foci numbers are shown separately for EdU-negative (E-) and EdU-positive (E+) cells. Red lines are medians. P values were calculated in Wilcoxon tests. C) Magnitudes of differences between RPA32 S33P/EdU PLA foci distributions in STING WT vs. control and STING S358A vs. control cells were expressed as Cliff’s delta values and plotted. Shown are Cliff’s delta values for each of eight independent experiments performed and quantified as in (B). Each experiment is identifiable by fill tone or fill pattern of the circle symbols. Triangle symbols are means. Blue arrowheads identify the values from the experiment shown in (B). D) Magnitudes of differences between mean fluorescence intensities (MFI) per nucleus measured by IF in situ for RPA32 S33P in STING WT vs. control and STING S358A vs. control cells were expressed as Cliff’s delta values and plotted. MFI values were measured in EdU-positive, HU-arrested cells. In (B-D) cells were labeled with EdU for 20 min and arrested with 5mM HU for 6 hrs in no-EdU media.

